# Secretome analysis of oligodendrocytes and precursors reveals their roles as contributors to the extracellular matrix and potential regulators of inflammation

**DOI:** 10.1101/2024.07.22.604699

**Authors:** Marlesa I. Godoy, Vijaya Pandey, James A. Wohlschlegel, Ye Zhang

## Abstract

Oligodendrocytes form myelin that ensheaths axons and accelerates the speed of action potential propagation. Oligodendrocyte progenitor cells (OPCs) proliferate and replenish oligodendrocytes. While the myelin-forming role of oligodendrocytes and OPCs is well-established, potential additional roles of these cells are yet to be fully explored. Here, we analyzed the secreted proteome of oligodendrocytes and OPCs *in vitro* to determine whether these cell types are major sources of secreted proteins in the central nervous system (CNS). Interestingly, we found that both oligodendrocytes and OPCs secret various extracellular matrix proteins. Considering the critical role of neuroinflammation in neurological disorders, we evaluated the responses and potential contributions of oligodendrocytes and OPCs to this process. By characterizing the secreted proteomes of these cells after pro-inflammatory cytokine treatment, we discovered the secretion of immunoregulators such as C2 and B2m. This finding sheds new light on the hitherto underappreciated role of oligodendrocytes and OPCs in actively modulating neuroinflammation. Our study provides a comprehensive and unbiased proteomic dataset of proteins secreted by oligodendrocyte and OPC under both physiological and inflammatory conditions. It revealed the potential of these cells to secrete matrix and signaling molecules, highlighting their multifaceted function beyond their conventional myelin-forming roles.

## Introduction

OPCs and oligodendrocytes are a significant cell population in the brain, comprising around 22-28% of the total cells [1, 2] Oligodendrocytes are responsible for generating myelin, a protective layer that ensheaths axons, expediting the propagation of action potentials and playing an essential role in axonal metabolism and health. In both developing and adult brains, oligodendrocyte progenitor cells (OPCs) undergo proliferation and differentiation to produce and maintain a population of oligodendrocytes.

While the myelin forming function of these cells are widely recognized, the possibility of oligodendrocytes and OPCs having other functions remains less understood. OPCs are distributed across both white and gray matter in the adult brain and account for more than 70% of dividing cells, making them the largest population of dividing cells in the adult brain [3]. However, the rate of differentiation of OPCs into mature myelinating oligodendrocytes decreases with age, leaving a large fraction of OPCs in the progenitor state throughout adulthood [3]. If only a small portion of the population is needed to replenish mature oligodendrocytes, why does the brain require a large number of OPCs to be present throughout adulthood?

OPCs are unique among the glial population in that they can form postsynaptic contact with neurons allowing them integrate neuronal activity with high spatial and temporal precision through the detection of quantal levels of neurotransmitter release [4]. However, the function of these synaptic connections and the effects of neuronal activity on OPCs is yet to be well understood.

Other major glial cells, such as microglia and astrocytes are key sources of signaling molecules [5, 6]. However, whether oligodendrocytes and OPCs also serve as significant contributors to the production of secreted signaling proteins remains unclear. Proteomics can characterize the landscape of secreted proteins from purified cell cultures. While the secreted proteome of oligodendrocytes has previously been profiled [7, 8], a similar characterization for OPCs has yet to be reported to our knowledge.

Neuroinflammation is a critical process in the regulation of CNS homeostasis and the resolution of pathological processes. However, when neuroinflammation is persistent or uncontrolled, it can contribute to the development of various neurodegenerative disorders, including Alzheimer’s disease, Parkinson’s disease, multiple sclerosis [9, 10]. This makes understanding the underlying cellular mechanisms of chronic or uncontrolled neuroinflammation crucial for developing new therapeutic strategies for these debilitating conditions.

While other glial cells, such as astrocytes and microglia, modulate neuroinflammation, whether OPCs and oligodendrocyte actively contribute to neuroinflammation is less understood. Interestingly, transcriptome changes in OPCs and oligodendrocytes have been reported in several neurodegenerative disease, such as Alzheimer’s disease and Parkinson’s disease, in addition to demyelinating diseases such as multiple sclerosis. A previous study suggested that OPCs can contribute to inflammation by participating in the breakdown of the blood brain barrier through interference of astrocyte endfeet and tight junctions, thus allowing the infiltration of inflammatory cells [4]. Similarly, OPCs migrate to the site of axonal injury and contribute to glial scar formation that consequently prevents regeneration of the damaged axons through the upregulation of genes for encoding for chondroitin sulfate proteoglycans (CSPGs) [4]. These observations highlight an urgent need to better understand the changes of OPCs and oligodendrocytes undergo in neuroinflammation and whether these changes actively contribute to the initiation, maintenance, and/or resolution of neuroinflammation.

In this study, we used an *in vitro* culture system to proliferate and differentiate OPCs and oligodendrocytes in both homeostatic and inflammatory conditions. Using a proteomics approach, we analyzed the secreted proteome of oligodendrocytes and OPCs, revealing their secretion of dozens of proteins, including various extracellular matrix proteins. We further examined their responses to pro-inflammatory cytokine treatment, discovering their secretion of immunoregulators such as C2 and B2m, thus illuminating their potential role in modulating neuroinflammation. Our comprehensive proteomic dataset highlights the potential of these cells to secrete matrix and signaling molecules, suggesting new functions of oligodendrocytes and OPCs beyond their well-known myelin-forming function.

## Materials and Methods

### Experimental animals

All animal experimental procedures were approved by the Chancellor’s Animal Research Committee at the University of California, Los Angeles, and conducted in compliance with national and state laws and policies. Sprague Dawley rats (dam with pups) were purchased from Charles River Laboratories and housed in standard cages. Rooms were maintained on a 12-hour light/dark cycle.

### OPC purification and culture

Whole brains excluding the olfactory bulbs from two pups at postnatal day 7 to day 8 were used to make each batch of OPC culture. OPCs were purified using an immunopanning method described before [11]. Briefly, the brains were digested into single-cell suspensions using 12u/ml papain. Astrocytes and differentiated oligodendrocytes were depleted using mouse anti-human integrin beta-5 (ITGB5) (eBioscience, 14-0497-82), Ran2 and galactocerebroside (GalC) hybridoma-coated panning plates, respectively. OPCs were then collected using an O4 hybridoma-coated panning plate. Cells were plated on 24-well plates at a density of 100,000 per well for secretome analysis and at a density of 20,000 cells per well for immunocytochemistry. For all experiments, OPCs were kept in serum-free proliferation medium containing growth factors PDGF (10 ng/ml, Peprotech,100-13A), CNTF (10 ng/ml, Peprotech, 450-13), and NT-3 (1 ng/ml, Peprotech, 450-03) for two days as previously described [11]. For oligodendrocyte differentiation, OPCs were switched to differentiation medium containing thyroid hormone (40 ng/ml, Sigma,T6397-100MG) but without PDGF or NT-3 for two days to differentiate them into oligodendrocytes as previously described [11]. Half of the culture media was replaced with fresh media every other day. All the cells were maintained in a humidified 37°C incubator with 10% CO_2_. Cells from both female and male rats were used.

### Cytokine treatment

Stock solutions at a concentration of 50μg/mL diluted in 1x PBS were made for TNF-α (Sigma, T5944-10UG) and Interferon-γ (Peprotech, 400-20) and stored at -80°C until use. TNF-α was added at a final concentration of 15ng/mL. Interferon-γ was added at a final concentration of 300ng/mL. OPCs were treated with cytokines at 2 days in vitro (DIV 2). Oligodendrocytes were treated with cytokines at 2 days post differentiation. Cells were incubated for 6 hours, 48 hours, or 5 days after treatment was added in a humidified 37°C incubator with 10% CO2.

### Media collection and trichloroacetic acid (TCA) precipitation

500μL of media was collected and spun at 200xg for 15min to remove dead cell debris. 400μL of the supernatant was removed and added to a new tube and 100% TCA (Sigma. T0699-100 mL) was added to a final concentration of 20%. Tubes were incubated on ice for 2 hours at 4°C. Tubes were then spun in a refrigerated microcentrifuge at 21.1xg for 15 min. Supernatant was removed leaving behind the protein pellet. Pellet was washed with 500μL of ice-cold acetone (FisherSci, A18-500) and spun at 21.1xg for 15 min twice. Acetone was removed after the last wash and pellet allowed to air-dry in fume hood until all residual acetone was removed. Pellets were parafilmed and stored at 80°C until used.

### Immunocytochemistry

Cells were washed with 1x PBS one time and fixed with 4% PFA (FisherSci, 50-980-495) for 15min at room temperate. Cells were then washed three times with 1x PBS for 5min. Cells were then blocked in a solution of 1x PBS+10% Normal donkey serum and 0.2% Triton-x100 (blocking solution) for 1 hour at room temperature. Next, antibodies against Pdgfr-α (1:100, R&D systems, AF1062) for OPCs, or Gal-c (1:5 of hybridoma supernatant) for oligodendrocytes, and NF-κB p65 (1:200, Cell Signaling, 8242S) were diluted in the blocking solution and added to the cells and incubated at 4°C overnight. The following day cells were washed three times with 1x PBS for 5min and incubated with fluorescent secondary antibodies (1:500, Invitrogen) for 1 hour at room temperature. Lastly, cells were washed three times in 1x PBS for 5min and mounted on glass slides (12-550-15, Fisher Scientific) using Dapi-Fluoromount-G (FisherSci, OB010020) and stored at -20°C until imaged. Coverslips were imaged at 20x on a Carl Zeiss Apotome epiflorescent microscope.

### Liquid Chromatography tandem mass spectrometry (LC/MS-MS)

TCA precipitated protein pellets isolated from conditioned media of control and treated oligodendrocytes and oligodendrocyte precursor cells were resuspended in 8M urea, 100mM Tris-Cl, pH 8.0 followed by reduction and alkylation by the sequential addition of 5 mM tris(2-carboxyethyl)phosphine and 10 mM iodoacetamide. This was followed by treatment with single-pot, solid-phase-enhanced sample preparation (SP3) protocol for protein clean-up [12]. Following SP3, eluates were proteolytically digested with Lys-C and trypsin at 37°C overnight. The digested peptides were subjected to offline SP3-based peptide clean-up and subsequently analyzed by LC-MS/MS. Briefly, peptides were separated by reversed phase chromatography using 75 μm inner diameter fritted fused silica capillary column packed in-house to a length of 25 cm with bulk 1.9μM ReproSil-Pur beads with 120 Å pores. The increasing gradient of acetonitrile was delivered by a Dionex Ultimate 3000 (Thermo Scientific) at a flow rate of 200nL/min. The MS/MS spectra were collected using data dependent acquisition on Orbitrap Fusion Lumos Tribrid mass spectrometer (Thermo Fisher Scientific) with an MS1 resolution (r) of 120,000 followed by sequential MS2 scans at a resolution (r) of 15,000. The data generated by LC-MS/MS were analyzed on MaxQuant bioinformatic pipeline [13]. A minimum of 2 peptides were selected for quantification. The Andromeda integrated in MaxQuant was employed as the peptide search engine and the data were searched against Rattus norvegicus (Uniprot Reference UP000002494). Briefly, a maximum of two missed cleavages was allowed. The maximum false discovery rate for peptide and protein was specified as 0.01 both at the level of peptide spectrum match and protein at the stage of identification. Label-free quantification (LFQ) was enabled with LFQ minimum ratio count of 1. The parent and peptide ion search tolerances were set as 20 and 4.5 ppm respectively. The MaxQuant output files were subsequently processed for statistical analysis of differentially enriched proteins using Analytical R tools for mass spectrometry (artMS) [14].

### Quantification and statistical analysis

#### Quantification of NF-κB p65

quantification of images was preformed when the experimenter is blinded to the treatment conditions on Fiji [15]using the cell counter function. Data comparisons between untreated and TNF-α/ IFN-γ treated were conducted with the Prism 9 software (Graphpad) using an unpaired two-tailed Welch’s T-test. An estimate of variation in each group is indicated by the standard error of the mean (S.E.M.) with * p<0.05, **p<0.01.

#### Proteomics dataset analysis

For all downstream analysis intensity measures were used. A protein list comprised of Log_2_ intensity values for the entire dataset was filtered using the following criteria before running statistical tests: 1) intracellular proteins were removed by comparing detected protein against a list of known rat secreted proteins obtained from the UniPort Consortium [16] to obtain a list of secreted proteins. 2) To remove potential contamination from astrocytes all proteins whose gene expression was 10-fold higher in astrocytes compared to OPC or oligodendrocytes were removed using gene expression data obtained from our previous work [17]. 3) the list of remaining proteins was then subdivided into an OPC and an oligodendrocyte list and proteins with a value of 0 intensity across all collected samples for their respective cell-type were removed. After filtering, the final dataset underwent quantile normalization, and the normalized dataset was used for statistical analysis. This generated a final list of 73 OPC-secreted proteins and 51 oligodendrocyte-secreted proteins.

#### Timepoint comparisons

T-tests with multiple comparison correction using the Two-stage step-up (Benjamini, Krieger, and Yekutieli) method were run for each cell type at each of the collection timepoints to compare differences between untreated and TNF-α/IFN-γ treated conditions using Prism 9 software (Graphpad).

#### Volcano plots

For each time point a dataset of proteins with detected intensity in at least 1 sample were compiled for each cell type. The average values of intensity for untreated and cytokine-treated were calculated for each timepoint for both cell types. The averages were used to calculate fold change which was transformed into a Log_2_(fold change) value. P-values were calculated using a t-test and then an adjusted p-value (Padj) was calculated using the benjamini Hochberg FDR correction (https://www.sdmproject.com/utilities/?show=FDR). The negative Log_10_(Padj) was then calculated. Proteins were classified as “upregulated” if they had a Log_2_ fold change greater than or equal to 1.5 and a Padj value of <0.05. Proteins were classified as “downregulated” if they had a Log_2_ fold change less than or equal to -1.5 and a Padj value of <0.05. Volcano plots were generated on R using the tidyverse package [18]

#### Heatmaps

Using the final list of 73 OPC-secreted proteins and 51 oligodendrocyte-secreted proteins, the average intensity value for the untreated samples at 48 hours and 5 days for each cell type was calculated. The protein list for each cell typed was rank based on this new average value and the top-10 detected proteins were graphed using the pheatmap (RRID:SCR_016418) package in R.

#### Venn diagram

For the comparison of shared and unique proteins between OPCs and oligodendrocytes, a dataset of normalized intensity values for only the untreated samples (8 for OPCs and 9 for oligodendrocytes) was used. Only proteins present in 3 or more samples of the same cell type were used for the comparison; proteins present in both cell types were labeled as “shared” and proteins present in only one of cell types were labelled as “unique”.

#### String dataset

The protein-protein interaction networks function enrichment analysis was performed on STRING [19]. For OPCs and oligodendrocytes a dataset of normalized intensity values for proteins present in only the untreated samples was used. The protein list was run on STRING and compared against a background list of all genes expressed by both OPCs and oligodendrocytes populations.

## Results

### Immunopanning purification and culture of rat OPCs and oligodendrocytes

To establish highly enriched OPC and oligodendrocyte cultures for our proteomic experiments, we used the immunopanning technique previously described in [11]. Briefly, we harvested whole rat brains, excluding the olfactory bulbs, at postnatal days 6-8, dissociated brain tissue into a single cell suspension, and depleted astrocytes, microglia, and oligodendrocytes through sequential antibody binding-based removal. OPC were collected by letting the single cell suspension bind to a petri dish coated with anti-O4 hybridoma, which binds OPCs (Figure 1A). To maintain OPC cultures, growth factors were included in the culture medium to promote proliferation and inhibit differentiation. To obtain oligodendrocyte cultures, we removed the growth factors, which led to the differentiation of OPCs into oligodendrocytes. To confirm the purity of our cultures we stained with antibodies against PDGFr-α and galactocerebroside (Galc), markers of OPC and oligodendrocytes respectively (Figure 1B) and confirmed more than 90% purity for the selected cell type.

**Figure 1:**
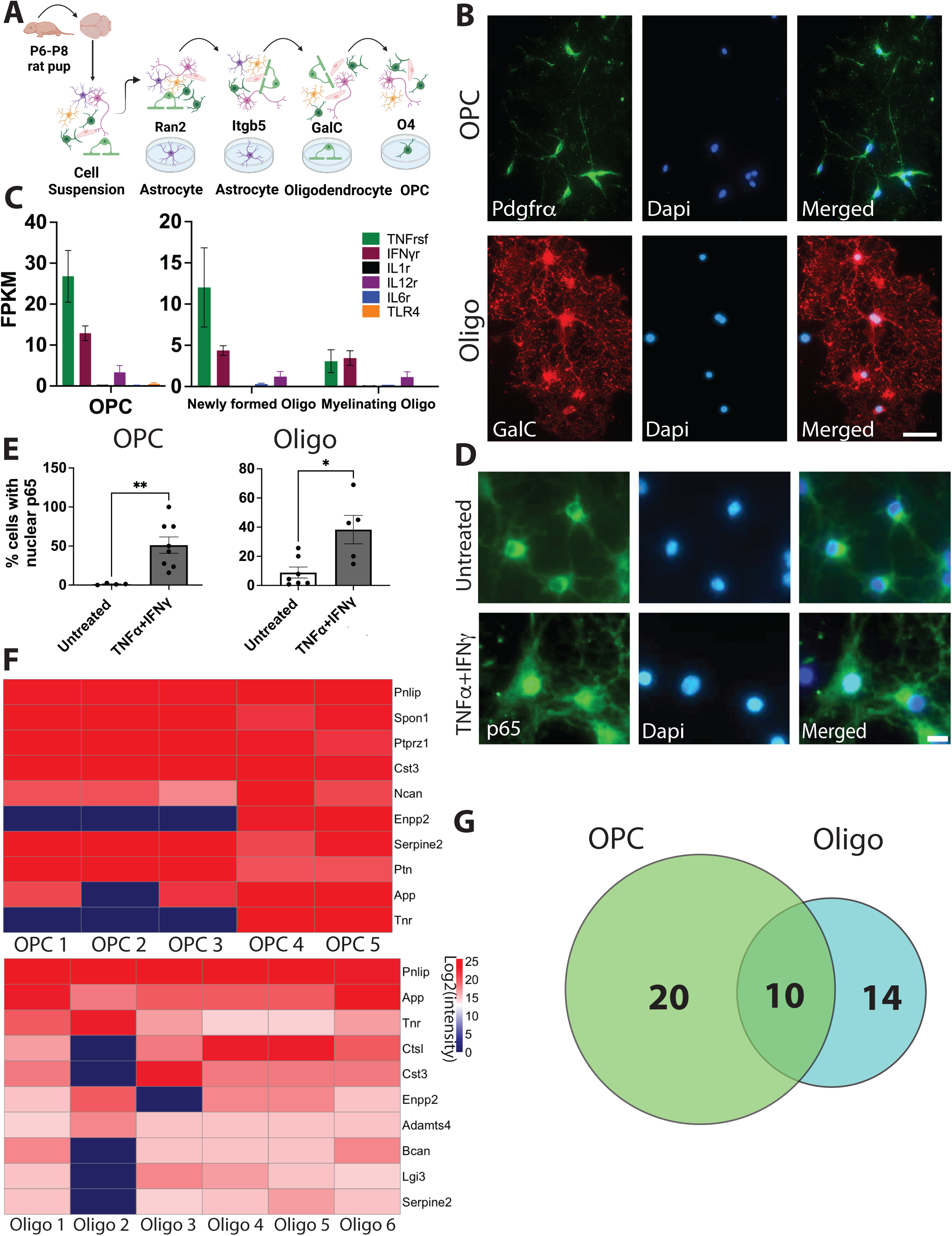
Purified rat Oligodendrocytes precursor cells (OPCs) and oligodendrocytes can be activated by TNF-α and IFN-γ. (A) Schematic of enzymatic digestion of rat whole brain to a single cell suspension followed by immunopanning purification using O4 hybridoma to bind oligodendrocyte progenitor cells (OPCs). (B) Representative images of cell culture purity following immunopanning procedure, OPCs (top) identified as PDGFrα positive (green) and oligodendrocytes (bottom) identified as galactocerebroside (GalC) positive (red); scale bar=50μm. (C) Expression of cytokine receptors in the oligodendrocyte lineage determined by RNAseq expression (Zhang *et al* 2014). (D) Representative images of p65 immunostaining in untreated (top) and TNF-α and IFN-γ treated (bottom) cells; scale bar =10μm. (E) quantification of nuclear p65 in untreated vs TNF-α and IFN-γ treated OPCs (left) and oligodendrocytes (right); two-tailed unpaired Welch’s T-test. Data are mean ± SEM, datapoints are number of images counted; For OPCs: untreated N=745 cells, treated N=293 cells, p=0.002; For oligodendrocyte: untreated N=720 cells, treated N=306 cells, p=0.0357. *p≤0.05, **p≤0.01 (F) Heatmap of the top-10 secreted proteins from untreated OPCs (top) and untreated oligodendrocytes (bottom); scale is Log2(intensity) from proteomics analysis. (G) Venn diagram of unique and shared proteins in untreated OPCs and oligodendrocytes generated using a dataset of normalized intensity values for only the untreated samples (8 for OPCs and 9 for oligodendrocytes). Only proteins present in 3 or more samples of the same cell type were used for the comparison.

### OPCs and oligodendrocytes secrete proteins associated with extracellular matrix organization

To define the secreted proteome of oligodendrocytes and OPCs, we analyzed the culture media of OPCs and oligodendrocytes by liquid chromatography tandem mass spectrometry (LC-MS/MS) and label-free quantitation (Figure 2C). This approach identified 73 and 51 secreted proteins for OPCs and oligodendrocytes, respectively (Supplementary Table 1 and Figure 1F).

**Figure 2:**
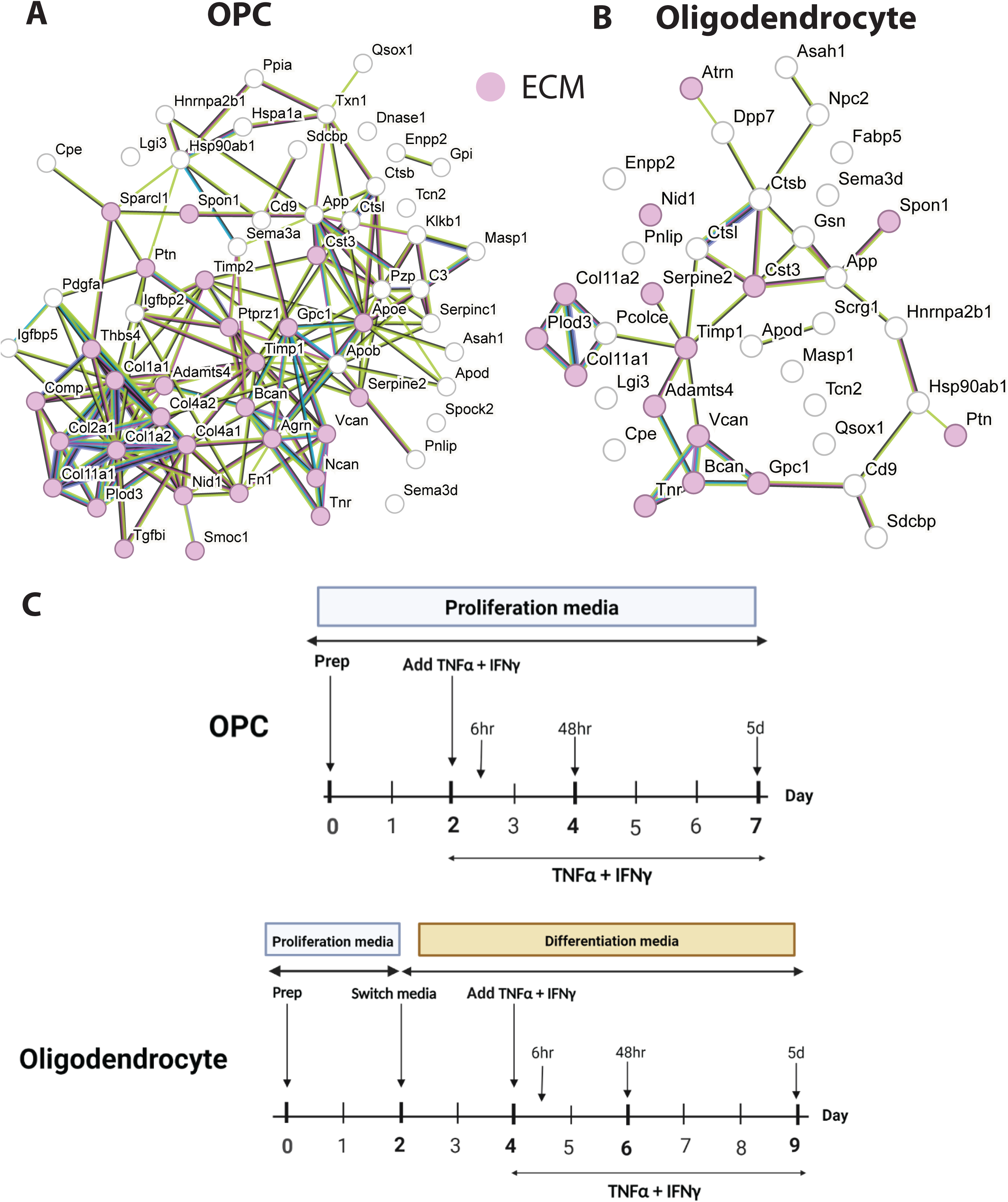
OPCs and oligodendrocytes secrete ECM associated proteins. STRING protein-protein network enrichment analysis of untreated OPC (A) and oligodendrocyte (B) secreted proteins. Red=associated with GO TERM “extracellular matrix” with FDR ≤0.05. (C) schematic of experimental timeline depicting when TNF-α and IFN-γ was added and when samples were collected.

We performed STRING protein-protein network enrichment analysis and detected enrichment of the gene ontology term for extracellular matrix (ECM) among both oligodendrocyte and OPC-secreted proteins (Figure 2 A-B, Supplementary Table 3). Both OPCs and oligodendrocytes secrete ECM proteins such as ECM structural components collagen, fibronectin, chondroitin sulfate proteoglycans (CSPGs, e.g., Neurocan, Versican, and Brevican) as well as proteases that may degrade and remodel the ECM (e.g., A Disintegrin And Metalloproteinase With Thrombospondin Motifs 4 (Adamts4), Serpin Family E Member 2 (Serpine2), Cathepsin L (Cstl) and Cystatin-C (Cst3)) (Fig. 2 and Supplementary Table 1). Of particular interest is the secretion of protease Adamts4, which degrades ECM component Aggrecan. The levels of aggrecan present in the perineuronal nets surrounding synapses is associated with restriction of neural plasticity and learning [20]. These results highlight ECM deposition and remodeling as a potential role of oligodendrocytes and OPCs in the brain as well as suggesting that OPCs and oligodendrocytes may have additional roles in modulating learning and memory by changing the structure of the ECM through protease secretion.

Comparing the secretomes of OPCs and oligodendrocytes we identified common proteins secreted by both cell types (e.g., Serpine2, Pancreatic Lipase (Pnlip), Cystatin C (Cst3), Ectonucleotide Pyrophosphatase/Phosphodiesterase 2 (Enpp2), Amyloid Beta Precursor Protein (APP), Tenascin R (Tnr), Spondin 1 (Spon1) suggesting OPCs and oligodendrocytes may have some shared roles in ECM maintenance (Tnr, Spon1) and neurite and axonal guidance (App, Enpp2, Serpine2) by secreting shared proteins into their environment. We also found there were a number of OPC-specific secreted proteins such as ECM-related molecules Neurocan (Ncan), SPARC-Related Modular Calcium-Binding Protein 1 (Smoc1), Fibronectin 1 (Fn1), Cartilage Oligomeric Matrix Protein (Comp), Collagen, Type III, Alpha 1 (Col3a1), Collagen, Type I, Alpha 1 (Col1a1), and Collagen, Type I, Alpha 2 (Col1a2) suggesting OPCs play a role in the structural integrity of the ECM. Likewise, we also found a few oligodendrocyte-specific secreted proteins such as Semaphorin 3D (Sema3d), Fatty Acid Binding Protein 5 (Fabp5), N-Acylsphingosine Amidohydrolase 1 (Asah1), and Mannose-Binding Lectin-Associated Serine Protease 1 (Masp-1), a component of the lectin pathway of complement activation [21, 22]. This suggests oligodendrocytes may have roles in axonal guidance (Sema3d) and signaling the innate immune system (Fabp5, Asah1) and the complement pathway (Masp1) (Figure 1F, G and Supplementary Table 2).

### TNF-α and IFN-γ treatment induces NFκB activation in OPCs and oligodendrocytes

Having characterized the secretomes of OPCs and oligodendrocytes in healthy cultures, we next investigated how their secretomes may change in neuroinflammation. Cytokines are crucial signaling molecules that induce reactive responses of microglia, astrocytes, and recruit and activate peripheral immune cells. To identify the cytokines to which OPCs and oligodendrocytes may respond during neuroinflammation, we examined the expression of cytokine receptor genes to find the most highly expressed ones in the oligodendrocyte lineage. Using RNAseq data from a previous study [17], we plotted the expression levels of six receptor families and found that both TNF receptor superfamily and interferon-gamma receptor family were among the most highly expressed receptors in the oligodendrocyte lineage (Figure 1C).

To determine if TNF-α and IFN-γ could induce reactive responses in OPCs and oligodendrocytes, we used Nuclear-Factor κB (NFκB) activation as a readout. NFκB is a transcription factor well characterized for its control of genes involved in inflammatory responses [23] and is activated in reactive microglia and astrocytes. We used an antibody against the phosphorylated Rel/p65 (p65) subunit of the NFκB complex, which is phosphorylated and translocates from the cytoplasm to the nucleus upon activation of the NFκB pathway [24]. We treated both OPCs and oligodendrocyte cultures with a combination of TNF-α and IFN-γ for 48 hours and then stained the cells using antibodies against p65, together with cell type markers PDGFr-α and Gal-c, for OPCs and oligodendrocytes respectively (Figure 1D). After TNF-α and IFN-γ exposure, the percentage of OPCs and oligodendrocytes with nuclear p65 significantly increased (Figure 1E). This finding shows that both OPCs and oligodendrocytes are responsive to the pro-inflammatory cytokines TNF-α and IFN-γ.

### OPCs and oligodendrocytes secreted inflammation associated proteins in response to TNF-α and IFN-γ treatment

To understand what proteins TNF-α and IFN-γ-treated OPCs and oligodendrocytes secrete into their microenvironment and if their secretion profiles differ at distinct timepoints following an inflammatory stimulus, we treated cells with TNF-α and IFN-γ and collected conditioned media at 6 hours, 48 hours, or 5 days after the addition of cytokines into the media (Figure 2C).

In oligodendrocyte cultures we found cytokine-induced changes at all three timepoints collected. At the acute timepoint of 6 hours post treatment, we only saw an upregulation of secretion of pancreatic triglyceride lipase (Pnlip), a lipase protein that has been shown to be transiently upregulated in astrocytes following traumatic brain injury [25] (Figure 3A). At the intermediate timepoint of 48 hours, we saw changes in proteins that may regulate the inflammatory microenvironment, including a downregulation of metalloproteinase, Adamts4, and upregulation of inflammation associated proteins complement component C2 (C2), beta-2 microglobulin (B2M), C-X-C Motif Chemokine Ligand 9 (Cxcl9) and Procollagen C-endopeptidase enhancer-1 (Pcolce) (Figure 3B). C2 is part of the classical pathway of complement activation and plays a role in innate immune response as a serine protease. B2M is a component of the class 1 major histocompatibility complex which is involved in antigen presentation to the immune system. *In vivo* B2M expression allows oligodendrocyte antigen presentation to CD8+ T-cells and consequently recruit the adaptive immune response [26]. Adamts4 is a key enzyme in the degradation of CSPGs, which are upregulated at the glial scar after CNS injuries and exhibit both beneficial and detrimental effects [27], including augmenting the pro-inflammatory responses [28]. Similarly, Pcolce, a metalloproteinase inhibitor, is a key regulator of the fibrillogenesis of collagen, a component of the glial scar [29]. These observations suggest that oligodendrocytes may regulate glial scar formation through secreted proteins. The upregulated secretion of inflammatory associated proteins and regulators of glial scar formation supports the idea that at 48 hours post-TNF-α and IFN-γ, oligodendrocytes are actively playing a role in modulating the inflammatory microenvironment in the brain.

**Figure 3:**
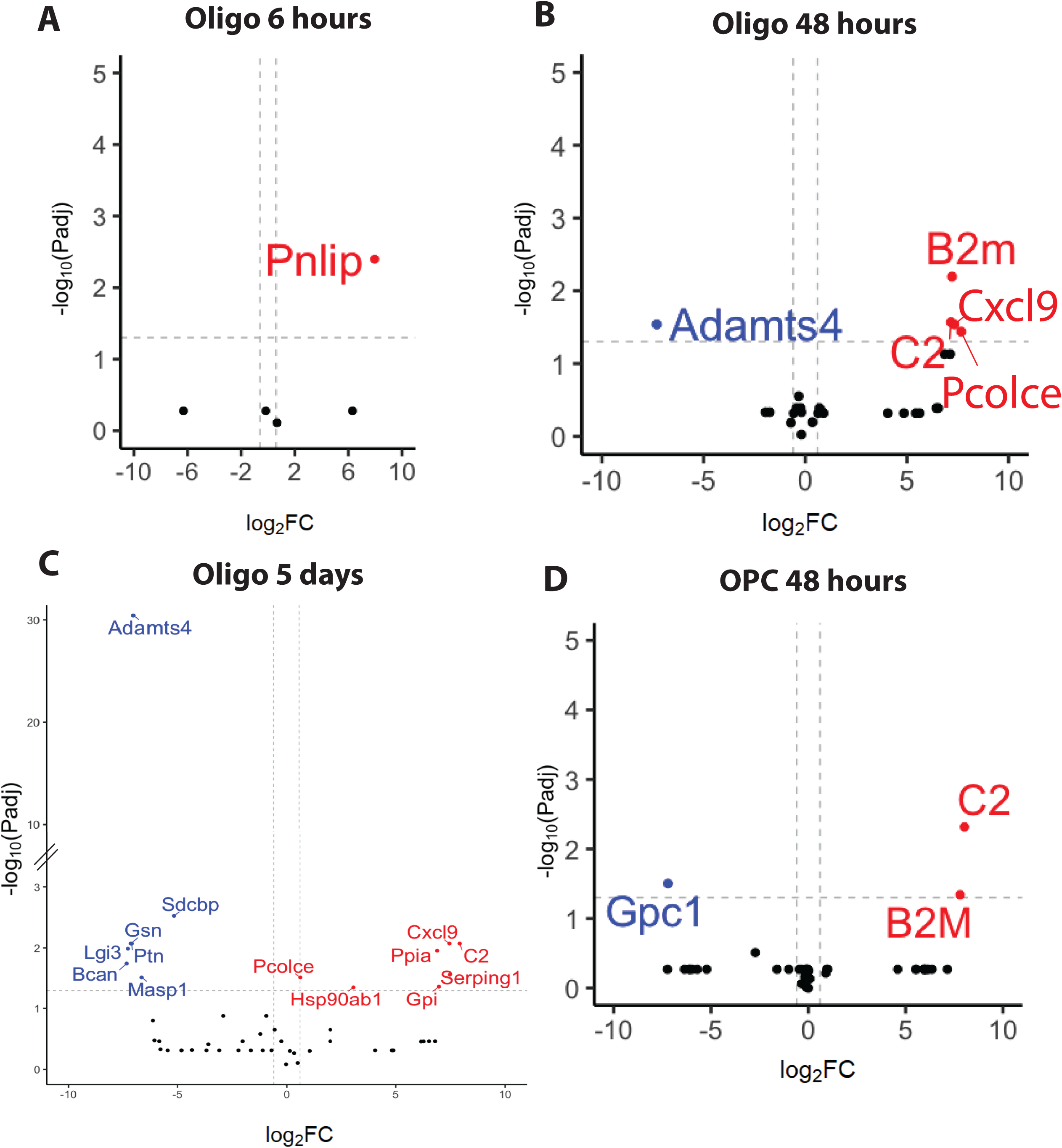
The secretomes of OPCs and oligodendrocytes are altered after treatment with TNF-α and IFN-γ. (A-D) volcano plots of the differential protein expression analysis for untreated versus TNF-α and IFN-γ treated oligodendrocytes at 6 hours (A), 48 hours (B), and 5 days after the start of treatment (C), and OPCs at 48 hours (D). Upregulated proteins are depicted in Red (criteria: fold change ≥ 1.5 and Padj ≤0.05). Downregulated proteins are depicted in Blue (criteria: fold change ≤ 0.67 and Padj ≤0.05). Dashed lines are drawn at fold change = 1.5 and Padj=0.05.

In contrast to the small number of changes at acute time points, at the late timepoint of 5 days post TNF-α and IFN-γ treatment, we saw a larger number of proteins (7 upregulated, 6 downregulated) significantly change after cytokine exposure. Oligodendrocytes downregulated secretion of inflammatory response associated proteins such as gelsolin (Gsn), an actin depolymerizing protein describe has having a modulatory role in inflammation [30], Masp-1, an activator of the complement pathway, Brevican (Bcan), a CSPG which has been shown to be secreted after brain injury by reactive astrocytes [31], and pleiotrophin (Ptn), a trophic factor upregulated in diseases characterized by neuroinflammation such as Alzheimer’s disease and Parkinson’s disease [32, 33] (Figure 3C). Interestingly, secretion of proteins associated with anti-inflammatory properties such as Serpin Family G Member 1 (Serping1) [34, 35], Heat Shock Protein 90 Alpha Family Class B Member 1 (Hsp90ab1) [36, 37], and neurotrophic factor neuroleukin (Gpi) [37–39] were upregulated at 5 days post cytokine treatment (Figure 3C). This combination of proteins suggests that at 5 days post injury oligodendrocytes may be playing a multi-faceted role in the modulation of neuroinflammation by contributing to both pro and anti-inflammatory mechanisms.

Together, our temporal analysis of protein secretion in oligodendrocytes following TNF-α and IFN-γ treatment reveals oligodendrocytes have a multi-phasic response to cytokines. Acutely at 6 hours, oligodendrocytes do not secrete inflammation associated proteins. At an intermediate timepoint of 48 hours after cytokine treatment, we began to see secretion of inflammation associated proteins such C2, B2M, and Cxcl9. At 5 days following cytokine treatment, we saw the continued secretion of C2 and Cxcl9 suggesting these two proteins may play a key role in the continued oligodendrocytes response to inflammation. Once oligodendrocytes are stimulated by cytokines, they display a dynamic and sustained response that modulates the immune response. Understanding the full range of the oligodendrocyte response to inflammation may garner new avenues of targeted treatments for conditions such as multiple sclerosis and Alzheimer’s disease.

On the other hand, in OPC cultures we found that treatment with TNF-α and IFN-γ did not significantly change protein secretion at the acute timepoint of 6 hours or the late timepoint of 5 days after the beginning of treatment. However, we did see a significant change after 48 hours of treatment in OPCs. TNF-α and IFN-γ exposure resulted in the downregulation of Glypican-1 (GPC-1) secretion and the upregulation of C2 and B2M secretion (Figure 3D). GPC-1 is a cell surface heparan sulfate proteoglycan that interact with a number of ligands to regulate numerous cellular functions and is overexpressed in certain cancers [38]. The secretion of C2 and B2M implies that OPCs, much like oligodendrocytes, are actively involved in regulating the inflammatory response. Together our findings suggests OPCs may play a role in modulating the inflammatory response following cytokine treatment.

In summary, our findings indicate functional roles for OPCs and oligodendrocytes beyond myelination. Both OPCs and oligodendrocytes secrete components of the ECM and proteases that suggest an active role in modulating the ECM in homeostatic conditions. In the presence of cytokines, OPCs and oligodendrocytes are able to contribute to neuroinflammation via secretion of immunoregulators such as C2, B2M, and Cxcl9. Our work also suggests that both OPCs and oligodendrocytes may function in temporally complex roles during an injury by changing the proteins they secrete. OPCs appear to respond to cytokines 48 hours after exposure by upregulating secretion of C2 and B2M. On the other hand, oligodendrocytes are more dynamic and respond to cytokines at acute, intermediate, and late phases after expose revealing potential transient responders to injury. These findings offer a useful unbiased proteomic dataset that exposes the multifunctional roles of OPCs and oligodendrocytes as contributors to the ECM and as potential regulators of inflammation.

## Discussion

Inflammation plays a major role in a number of neurodegenerative diseases and CNS injuries such as stroke, Alzheimer’s disease, multiple sclerosis, and Huntington’s disease [39, 40]. It is crucial to understand how oligodendrocytes and OPCs communicate with other cell types, such as microglia and astrocytes, and how these interactions shape their response to inflammatory stimuli. Microglia and astrocytes have been shown to interact with oligodendrocytes in several ways [41, 42]. For instance, microglia have been shown to promote OPC proliferation and differentiation in response to inflammation [43]. Moreover, astrocytes can secrete cytokines and chemokines that regulate the immune response and influence oligodendrocyte function [44].

In the context of inflammation, the secretion of protein such as C2, B2M, Masp1 and Cxcl9 suggest that OPCs and oligodendrocytes may influence inflammatory responses by communicating with other glial cells. C2 and Masp1 are components of the classical and lectin complement systems. C2 can bind to membrane bound C4 on astrocytes and produce C3. C3 serves as a “tag” for cell or debris elimination by activating the C3r receptor on macrophages and microglia and promoting phagocytosis or activating the terminal complement cascade and inducing cell lysis [45]. Cxcl9 functions via binding to its receptor Cxcr3 located on astrocytes and microglia. Cxcl9 secretion by astrocytes has been described as pro-inflammatory and leading to recruitment of monocytes and cytotoxic leukocytes [5]. In a cuprizone model of demyelination using a knockout of Cxcr3, both microglial and astrocyte reactivity was decreased following demyelination [46]. Likewise, Cxcr3 signaling in an APP/Ps1 mouse model of Alzheimer’s disease has been shown to increase plaque formation and reduce microglial phagocytosis of plaques [47]. Our discovery of Cxcl9 secretion by oligodendrocytes under inflammatory conditions points to a novel role of oligodendrocytes in manipulating the Cxcl9/Cxcr3 axis to regulate glial reactivity at the site of injury. Similarly, B2M is a component of the class 1 major histocompatibility complex which is involved in antigen presentation to the immune system. B2M in aging studies has been shown to induce microglial reactive responses and cause cognitive decline by increasing synaptic pruning [48] and reducing neurogenesis [49]. Secretion of B2M by oligodendrocytes and OPCs suggests that these cells may play a dynamic role in regulating the reactive responses of other glial cells and immune cells that either dampen or increase the inflammation following cytokine exposure. Microglia can release both TNFα and IFNγ in inflammatory conditions such as demyelination in multiple sclerosis, which may trigger the release of inflammatory signals for the oligodendrocytes [50]. It may be the case that both autocrine and paracrine signaling make OPCs and oligodendrocytes reactive and that reactive oligodendrocyte-lineage cells can secrete inflammatory signals that induce astrocytes and microglia reactivity. This finding reveals that targeting of OPCs and oligodendrocytes may modulate inflammation and represents a novel approach for treating inflammatory conditions such as Alzheimer’s disease and multiple sclerosis.

The ECM is a matrix comprised of glycoproteins, collagens, and glycosaminoglycans that have structural, signaling, and metabolic functions in the CNS [51]. Together these molecules can form basement membranes around blood vessels and the glia limitans, perineuronal nets around the soma of neurons, or the interstitial matrix between cellular structures [51, 52]. The ECM is also rich in chondroitin sulfate proteoglycans (CSPGs) that includes a family of lecticans (aggrecan, veriscan, brevican, and neurocan) that have quite extensively studied as components of the Node of Ranvier, playing a crucial role in action potential propagation. [51, 53].

CSPGs such and neurocan and brevican, inhibit the migration and differentiation of OPCs by creating a mechanical and chemical barrier hindering migration [54]. In addition, OPCs are mechanosensitive and highly susceptible to the stiffness of their surroundings; a softer matrix is more permissive to differentiation [52]. The secretion of proteases such as Adamts4, Serpine2, Cstl and Cst3 suggests that OPCs and oligodendrocytes may take an active role in regulating their microenvironment in unstimulated conditions. The expression of ECM molecules such as CSPGs, collagens, and fibronectins in conjugation with proteases could indicate that OPCs and oligodendrocytes play a role in ECM formation during development and structure maintenance in the homeostatic adult brain.

Our findings that OPCs and oligodendrocytes secrete ECM molecules and proteases supports previous findings by other groups that inflammatory cytokines such as TNF-α and IFN-γ can stimulate the release of ECM components, influence ECM turnover, and protease secretion [52, 55], while providing new evidence for the contribution of OPCs and oligodendrocytes to ECM deposition and remodeling. The ECM and its individual components and 3D ultrastructure can modulate immune function and responses by signaling cell migration or cell proliferation in or around the injury site [55]. CSPGs such as brevican and neurocan have been previous found to be secreted in response to injury and in vitro these molecules have been shown to inhibit axonal growth [56]. CSPGs are also main components of the glial scar [51] and multiple cell types including astrocytes, neurons, oligodendrocytes, and microglia have been known to contribute to ECM upregulation following injury [52]. In multiple sclerosis, the ECM is altered at the site of lesions and CSPGs such as neurocan have been shown to have enhanced expression at the edge of chronic lesions [52]. Recent proteome studies of multiple sclerosis plaques and brain and spinal cord of experimental autoimmune encephalomyelitis (EAE) mice showed an abundance of ECM components at active plaques and that ECM molecules were amongst the most highly expressed proteins in EAE [52]. ECM fragments can act as damage-associated molecular patterns (DAMPs) and interacting directly with toll-like receptors providing an active role in shaping the immune response [52]. Our study provided evidence that OPCs and oligodendrocytes may play a dynamic role in modulating the ECM in both homeostatic and inflammatory conditions.

Adamts4 is a protease abundant in the white matter and whose expression is restricted to mature oligodendrocytes [57] and is responsible for cleaving CSPGs such as brevican and neurocan [58]. Adamts4 protein levels are elevated in the EAE model of multiple sclerosis and spinal cord injury [59, 60]. Adamts4 cleaves brevican [61] which is the most abundant proteoglycan in the CNS and localizes near synapses [62]. Increases of ECM components brevican, neurocan and tenascin-r, which localize to perineuronal nets, were found to be increased in early mouse models of AD and correlated with a loss of contextual fear conditioning and LTP [59]. In addition, Adamts4 has been shown to cleave aggrecan, a CSPG and the backbone of the perineuronal nets that plays an important role in incorporation of other CSPGs, such as brevican, into the perineuronal structure; deletion of aggrecan removed perineural nets from parvalbumin interneurons and increased neuroplasticity [63, 64]. Similarly, Adamts4 also cleaves Reelin, a glycoprotein important for learning and memory, and an increased in cleaved Reelin leads to increased phosphorylation of Tau and formation of neurofibrillary tangles [59]. Secretion of Adamts4 at specific periods may play a role in learning and memory by cleaving lectins and loosening the extracellular matrix around neurons allowing for plasticity [62]. Together with these data, our finding of oligodendrocyte secretion of Adamts4 suggests that oligodendrocytes may play a role in modulating learning and memory through regulating the cleavage of ECM components.

Studying the alternative functions of OPCs has been a challenged faced by many in the field. The manipulation required to study OPC function often results in alterations to myelin and myelin formation, thus making it difficult to separate effects due to OPCs from the effects due to altered myelination [4]. Our study provides a way to infer alternative functions of OPCs and oligodendrocytes via their secreted proteins. The proteome of secreted proteins provides information on protein levels that transcriptomic studies cannot provide. Our findings support a functional role of OPCs and oligodendrocytes in regulating their microenvironment via secretion of ECM molecules and inflammatory proteins in both homeostatic and stimulated conditions. This study provides novel insight into potential mechanisms of communication between glial cells in the context of inflammation, learning and memory.

## Acknowledgements

This work is supported by the NIH/NIMH T32MH073526 and the Achievement Rewards for College Scientists Foundation Los Angeles Founder Chapter to M. I. G., the NIH/NINDS R00NS089780, R01NS109025, the NIH/NIA R03AG065772, NIH/NICHD P50HD103557, National Center for Advancing Translational Science UCLA CTSI Grant UL1TR001881, the BSCRC Innovation Award, the Friends of the Semel Institute for Neuroscience & Human Behavior Friends Scholar Award, the W.M. Keck Foundation Junior Faculty Award, UCLA Jonsson Comprehensive Cancer Center and BSCRC Ablon Scholars Award, Rose Hills Foundation Stem Cell Innovation Award to Y. Z.

## Supplementary Tables

**Supplementary Table 1**: Proteomics dataset including the raw, filtered, and normalized intensity values

**Supplementary Table 2:** List of genes used for unique and shared protein Venn diagram for OPCs and oligodendrocytes

**Supplementary Table 3**: List of genes for OPCs and oligodendrocytes used for STRING analysis and the Gene Ontology (GO) Terms for OPCs and oligodendrocytes generated by STRING

